# Metaproteogenomics resolution of a high-CO_2_ aquifer community suggests an active symbiotic lifestyle of groundwater Gracilibacteria

**DOI:** 10.1101/2023.12.18.572140

**Authors:** Perla Abigail Figueroa-Gonzalez, Till L. V. Bornemann, Tjorven Hinzke, Sandra Maaß, Anke Trautwein-Schult, Joern Starke, Carrie J. Moore, Sarah P. Esser, Julia Plewka, Tobias Hesse, Torsten C. Schmidt, Ulrich Schreiber, Batbileg Bor, Dörte Becher, Alexander J. Probst

**Author notes:** authors contributed equally (Perla Abigail Figueroa-Gonzalez, Till L. V. Bornemann).

## Abstract

**Background:** Bacteria of the Candidate Phyla Radiation (CPR), constituting about 25% of the bacterial biodiversity, are characterized by small cell size and patchy genomes without complete key metabolic pathways, suggesting a symbiotic lifestyle. Gracilibacteria (BD1-5), which are part of the CPR branch, possess alternate coded genomes and have not yet been cultivated. However, besides genomic evidence, little is known about the lifestyle of Gracilibacteria, their temporal dynamics, and activity in natural ecosystems, particularly in groundwater, where they were initially been genomically resolved. Therefore, we here aimed to investigate Gracilibacteria activity *in situ* and to discern expressed genes involved in their lifestyle, using the metaproteogenome of Gracilibacteria as a function of time in the cold-water geyser Wallender Born in the Volcanic Eifel region in Germany.

**Results:** We coupled genome-resolved metagenomics and metaproteomics to investigate a cold-water geyser microbial community enriched in Gracilibacteria across a 12-day time-series. Groundwater was collected and sequentially filtered to fraction CPR and other bacteria. Based on 670 Gbps of metagenomic data, 1129 different ribosomal protein S3 marker genes and 751 high-quality genomes (123 population genomes after dereplication), we identified dominant bacteria belonging to Galionellales and Gracilibacteria along with keystone microbes, which were low in genomic abundance but substantially contributing to proteomic abundance. Seven high-quality Gracilibacteria genomes showed typical limitations, such as limited amino acid or nucleotide synthesis, in their central metabolism but no co-occurrence with potential hosts. The genomes of these Gracilibacteria encoded for a high number of proteins related to a symbiotic or even predatory lifestyle, *e.g.*, type IV and type II secretion system subunits and features related to cell-cell interactions and cell motility, which were also detected on protein level.

**Conclusions:** Coupling metagenomics to metaproteomics enabled us to identify microbial keystone taxa in a high-CO_2_ aquifer, and to reveal microbial dynamics of Gracilibacteria. We posit that Gracilibacteria might be successful microbial predators in this ecosystem, potentially aiding in population control of this highly perturbed microbial geyser community from the deep biosphere.

## Background

Bacteria belonging to the Candidate Phyla Radiation (CPR, also known as Patescibacteria [1]) comprise at least one quarter of the bacterial domain [2,3], and are characterized by having reduced genomes as well as small cell sizes (<0.2 µm, [4–7]). In addition, CPR bacteria show a distinct differentiation in protein content and ribosomal structure when compared to other bacteria [4]. Due to the very few cultivated representatives of CPR bacteria, few biochemical studies on their proteins have been conducted resulting in many proteins of unknown function [8]. However, their core metabolic genes are limited and essential pathways for free-living bacteria, such as capabilities for *de novo* synthesis of lipids or amino acids or the TCA cycle, are usually very incomplete in CPR bacteria. Interestingly, genomes of CPR bacteria have been shown to encode a substantial amount of proteins with transmembrane helices, as well as proteins which are likely localized in the membrane or secreted [9]. With the increased use of metagenomic environmental surveys, CPR bacteria have been detected in a plethora of environments, including hydrothermal vents, soil, lakes, rivers, permafrost, the (deep) terrestrial subsurface and groundwater, as well as oral cavities of mammals [10–18]. Considering the limited metabolic potential and small genome size of CPR bacteria, which is comparable to symbiotic non-CPR bacteria [8], CPR bacteria likely depend on hosts in order to obtain essential molecular building blocks, such as amino acids, lipids, or co-factors [6,8]. This host-dependency was first shown to be the case for the phylum Saccharibacteria, and in particular for *Nanosynbacter lyticus* (also known as TM7x), which was isolated in co-culture with its host, *Schaalia odontolyticus* XH001, from the human oral cavity [15,19]. While *N. lyticus* can, however, achieve stable growth conditions along with its host [20–22], recent reports of predatory CPR bacteria, specifically of the Absconditabacteria lineage, show that the host is killed for the CPR bacteria to survive, regardless of the environmental conditions [23,24].

Gracilibacteria, originally named BD1-5 [7,25], is a relatively understudied phylum of CPR bacteria and forms a clade with the candidate phyla Abscondibacteria and Peregrinibacteria [23]. A trademark characteristic of the Gracilibacteria lineage, similar to Absconditabacteria, is their use of genetic code 25, where the codon UGA encodes for a glycine, as opposed to being a stop codon [26–28]. Like other members of CPR bacteria, Gracilibacteria have very limited metabolic capabilities (*e.g*., some members encode for the oxidative branch of the pentose pathway but lack steps for glycolysis and TCA). At the same time, Gracilibacteria encode for several surface proteins related to type IV pili and secretion systems [28], indicating that these organisms have the potential for cell-cell interactions.

One such example of predatory CPR bacteria is *Candidatus* Absconditicoccus praedator, isolated from a hypersaline alkaline lake along with its host *Halorhodospira halophila*, an obligately anaerobic photosynthetic purple-sulphur bacterium [24]. The symbiont’s metabolic limitations make it completely dependent on its host, whom it kills by lysing and consuming its cytoplasmic contents. A second case of a predatory CPR bacterium was reported for *Candidatus* Vampirococcus lugosii, discovered in an athalassic hypersaline lake in Spain. Cultures derived from the lake’s microbial mat showed that *Ca.* V. lugosii is an epibiotic bacterium that feeds on *Halochromatium*, an anoxygenic photosynthetic gammaproteobacterium [23]. Both of these CPR bacteria are members of the Absconditabacteria phylum. Similar to *Ca.* A. praedator, the attachment of *Ca*. V. lugosii leads to compromising the cell membrane of its host, so that *Ca*. V. lugosii can consume the *Halochromatium* bacterium’s cytoplasmic content, thereby killing it [23,24]. Based on *Ca.* A. praedator and *Ca*. V. lugosii’s predatory lifestyle, it was proposed that they might play a role in population control of their hosts. As there is as of yet no cultured Gracilibacteria representative, no information about their host range nor predatory character is available. In addition, while metabolic potential can be inferred based on genomes, the only proteomic report about Gracilibacteria focuses on their alternative genetic code and not on their protein expression, *i.e.*, actual phenotype, as for example related to metabolism or cell-cell interactions [7,26]. Consequently, whether they actually live as symbionts as predicted from their genomes, has remained elusive.

In this study, we characterized a model ecosystem (cold-water geyser Wallender Born (WB; 50°09’13.4”N 6°43’13.1”E)) enriched in Gracilibacteria, to investigate proteogenome of Gracilibacteria in a complex microbial community. The geyser is located in the town of Wallenborn, in the Volcanic Eifel region of Germany. The driving force for its eruptions is the accumulation of carbon dioxide (CO_2_) in groundwater. CO_2_ saturates the groundwater, which leads to the geyser’s eruption where groundwater and CO_2_ are released to the atmosphere. These eruptions occur in approximate 40 minute intervals. Taking advantage of these groundwater eruptions, we used geyser WB as a study site to characterize its microbial community on the molecular level. For this, we sampled erupted groundwater over a twelve-day time series and sequentially filtered it to enrich small- and large-sized microbes on 0.1-µm filters and 0.2-µm filters, respectively. We used genome-resolved metagenomics to obtain high-quality genomes, and coupled this to metaproteomics analyses of the same time points, to generate in-depth insights into Gracilibacteria metabolism and ecology. We identified traits related to a predatory lifestyle of Gracilibacteria in the geyser, both in their metabolic potential as well as in expressed proteins with metabolic similarities with *Ca.* A. praedator and *Ca*. V. lugosii. Ultimately, we discerned a predatory lifestyle of groundwater Gracilibacteria, hinting at their role as key members of the geyser’s microbial community.

## Material and methods

### Geological context of sampling site

The cold-water geyser Wallender Born (WB) is located in the town Wallenborn, southwest of the subrecently active Quaternary volcanic field of the West Eifel region, in the Rhenish Massif (50°09’13.4”N 6°43’13.1”E). It is located between the Manderscheid anticline to the southeast and the Salmtal syncline to the northwest which formed during the Variscan orogeny event. It consists of weakly metamorphic early Devonian (Emsian) sedimentary units [29]. From a geological point of view, it belongs to the eastern edge of the old depression zone of the Eifel north-south zone (Eifeler Nord-Südzone). This zone is characterized both by the Middle and Late Devonian Eifel limestone synclines, preserved in the depression, and by numerous N-S to NNE-SSW trending fault zones [30]. The strong CO_2_ outgassing is related to the magmatic activity of the West Eifel region, which together with the East Eifel volcanic field is attributed to a mantle plume underneath the Eifel (Eifel plume). Helium R/Ra values from the Wallender Born geyser show a clear mantle influx of ³He, implying magmatic intrusion [31]. Please refer to Supplementary Figure S1 for a scheme of the geyser.

### Groundwater sampling and geochemical analyses

Time-series sampling of biomass was done for twelve days, from the 14^th^ – 25^th^ of October 2020. Groundwater was collected in sterile, DNA-free containers during each eruption, for immediate filtering on-site. The groundwater was filtered sequentially on 0.2-µm and then 0.1-µm filters (see Supplementary Figure S2A for a scheme of the filtering setup). Before starting the filtration of each time-point, the system was rinsed with groundwater to get rid of stagnant water left over from the previous sampled time-point. Additionally, one bulk sample, filtered directly onto a 0.1-µm filter, was obtained. Approximately 100 L were filtered on each 0.2-µm filter, with the flow-throughs of two 0.2-µm filters being combined via a T-piece and filtered onto one 0.1-µm filter (please refer to Supplementary Table S1 for individual filtered volumes). Filters were then immediately transferred to a sterile falcon tube, stored on dry ice on-site, and later on transferred to −80° C until further processing. Temperature and pH were measured in triplicates with a thermometer and pH strips, respectively.

#### Total iron

100 µL of groundwater were mixed with 900 µL of 1 M HCl on-site. Samples were taken in triplicates and kept at 4°C till measurement. Total iron was measured by mixing of 100 µl of aliquot with 900 µl hydroxylamine, followed by addition of 80 µl reduced aliquot to 120 µl Ferrozine, incubation for 10 min in the dark, and photometric determination by a Tecan plate reader at 560 nm [32].

#### Ions

0.1-µm pore-size filtered water was used for ion measurements. Samples were measured in dilutions of 1:100 and 1:500 in distilled water to be able to accurately quantify both highly abundant ions and lower abundant ions. Samples were analyzed using the Dionex Aquion ion chromatography system (Thermo Scientific, USA). Anions were analyzed with a Dionex IonPac AG23-4 µm guard column, a Dionex IonPac AG23-4 µm 2 x 250 mm analytical column, as well as an AERS 500 Carbonate 2 mm suppressor, Ultimate 3000 heating element, and a DS6 heated conductivity cell detector. Cations were measured on a CS12A RFIC 2 x 250 mm analytical column and a CG12A RFIC detector. Measurements were done in technical triplicates.

#### Total organic carbon (TOC)

Unfiltered groundwater samples were stored at 4 °C until measurement. Prior to measurement, the samples were filtered through a 0.45-µm filter to remove large particles. 10 ml of sample were augmented with 0.5 ml 1 M HCl to remove residual carbonate present in cold-water geysers, verified via pH strips. For each time point, two biological samples were measured in triplicate on a TOC-5050 Analyzer with ASL-5000 (Shimadzu) and averaged after accounting for the HCl dilution.

### Epifluorescence microscopy

For each time-point, groundwater was filtered on-site through a 0.2-µm syringe-filter using a sterile syringe. The flow-through was collected in a sterile, DNA-free tube and subsequently filtered through a 0.1-µm syringe-filter using a second sterile syringe. Filters were incubated with 10 µg ml^-1^ DAPI in 2% (v/v) formaldehyde for five minutes, washed with distilled water, dried for ten minutes and stored in the dark at 4 °C until use. Cells were quantified using the Axio Imager M2m epifluorescence microscope equipped with an AxioCam MRm and a Zen 2 Pro software (Carl Zeiss Microscopy GmbH, Jena, Germany; version 2.0.0.0). Enumeration of the cells was done in ten horizontal and ten vertical fields, extrapolating counts to the entire filter area and normalizing through the filtered volume to calculate cells per millilitre.

### DNA extraction and sequencing

Each pair of 0.2-µm filters was pooled to have a comparable sample to the respective 0.1-µm filter (see Supplementary Figure S2B for a scheme of the pooling for metagenomics). This pooling resulted in 22 samples for the 0.2-µm filtered fractions, each corresponding to the different time points obtained during the sampling campaign, with their respective 0.1-µm filtered fraction. DNA extractions of the samples were performed using the Dneasy PowerMax Soil DNA Extraction kit (Qiagen, 12988-10) according to the manufacturer’s instructions and further concentrated using ethanol precipitation with glycogen as the carrier. Of the 44 samples, 30 yielded enough biomass for library preparation via the Westburg NGS DNA Library Prep Kit (cat. No. WB 9096). Of these, 28 were successfully sequenced using Illumina NextSeq500 (2x150 bp paired-end reads). Some libraries which yielded low sequencing depth in initial sequencing runs were sequenced up to a total of three times and the reads were concatenated to reach about 20 Gbps sequencing depth per sample.

### Sample preparation for metaproteomics

Due to the higher biomass requirements for metaproteomics as compared to metagenomics, sample sets analyzed differed from those for metagenomics. A total of three different metaproteomics sample sets were analyzed (see Supplementary Figure S2C and Supplementary Table S3 for details): I) We compiled a time-series consisting of individual 0.2-µm filters taken at different time points. II) To compare the proteome of the size-fractionated microbiomes, we separately pooled 0.2-µm filters and 0.1-µm filters of the respective time points to generate biomass consistent of only microbes from the 0.2-µm fraction and of the 0.1-µm fraction, respectively. This provided sufficient biomass for the 0.1-µm filter for most protein extraction but also generated one set of 0.2-µm filter samples without the corresponding 0.1-µm sample due to too-low biomass. III) For a whole-community, we used a bulk sample filtered directly onto a 0.1-µm filter (without size fractionation).

### Protein extraction

Protein extraction from the filters was performed according to previously published protocols [33] with some modifications. In brief, the filters were cut in pieces, transferred to low protein binding reaction tubes and covered with one volume of resuspension solution 1 (50 mM Tris-HCl pH 7.5, 0.1 mg ml^-1^ chloramphenicol, 1 mM PMSF (phenylmethanesulfonyl fluoride)). After vortexing, 1.5 volumes resuspension solution 2 (20 mM Tris-HCl pH 7.5, 2% (w/v) SDS) were added, and the samples were incubated at 60 °C for 10 min with vigorous shaking. After cooling down of the samples to room temperature, 5x volume of DNAse solution (1 μg ml^-1^ DNAse I) were added. Samples were lysed by ultrasonication on ice for 6 min (Sonopuls MS72 (Bandelin, Berlin, Germany), amplitude 51-60%; cycle 0.5), and subsequently incubated at 37°C for 10 min with vigorous shaking. After centrifugation, proteins were precipitated from the supernatants by adding pre-cooled trichloroacetic acid (TCA) to reach 20% (v/v) TCA and incubating at 4 °C for 30 min in an overhead inverter. Precipitates were pelleted and washed with pre-cooled acetone. The dried pellet was resuspended in 2x SDS sampling solution (0.125 M Tris-HCl pH 6.8, 4% (w/v) SDS, 20% (v/v) glycerol, 10% (v/v) ß-mercaptoethanol), and incubated in an ultrasonic bath for 15 min before heating at 95 °C for 5 min. Samples were centrifuged, supernatants were kept and the remaining pellet was again treated with 2x SDS sampling solution. Both sample fractions, supernatant and pellet, were loaded onto pre-cast SDS Gels (Criterion TGX 4-20%, 12+2 wells, Bio-Rad) and separated by SDS-PAGE. After staining with Coomassie, proteins were in-gel digested as described previously [34]. Briefly, gel lanes were excised in ten equidistant pieces and each piece was destained, washed, and subjected to trypsin digestion (as well as mass spectrometric measurement) individually. Peptides were eluted in water using an ultrasonic bath and desalted by the use of C18 ZipTip columns (Merck) according to the manufacturer’s guidelines.

### Metagenome processing

Illumina paired-end reads were checked for adapters and sequencing artefacts with Bbduk v37.09 (Bushnell, https://sourceforge.net/projects/bbtools), quality-trimmed with Sickle v1.33 (https://github.com/najoshi/sickle) and deduplicated with dedup (Bushnell, https://sourceforge.net/projects/bbtools). The diversity coverage in the samples based on raw reads was estimated using Nonpareil3 [35] in kmer mode with k=20 (*i.e.*, the minimal read length). For each individual metagenome, quality-controlled reads were first assembled with Metaviralspades 3.15.2 [36] to reconstruct viruses and plasmids, and paired reads where no read mate mapped to assembled viruses or plasmids (via Bowtie2 v2.3.5.1, –sensitive mode, [37]) were then assembled using metaspades 3.15.2 [38]. Viral and normal assemblies were then concatenated to yield the final assembly. Unless otherwise stated, all mappings were performed with Bowtie2 [37] in sensitive mode for, *e.g.*, determining relative scaffold abundances of the assembled scaffolds. Open reading frames were predicted on scaffolds with a minimum length of 1000 bps using Prodigal 2.6.3 [39] in meta-mode and annotated against FunTaxDB 1.3 [40] to retrieve function and taxonomic profiles.

### Mass spectrometry and data processing of metaproteomics samples

Peptides were subjected to LC-MS/MS analyses on a LTQ Orbitrap Elite instrument (ThermoFisher Scientific) coupled to an EASY-nLC 1200 liquid chromatography system. Therefore, peptides were loaded on a self-packed analytical column (OD 360 µm, ID 100 µm, length 20 cm) filled with 3 µm diameter C18 particles (Maisch) and eluted by a binary nonlinear gradient starting at 5% up to 99% acetonitrile in 0.1% (v/v) acetic acid over 82 min with a flow rate of 300 nL min^-1^. For MS analysis, a full scan in the Orbitrap with a resolution of 60,000 was followed by collision-induced dissociation (CID) of the 20 most abundant precursor ions. Database searches against a concatenated forward-reverse protein database derived from the described metagenome analyses (see “Generation of protein database from metagenomes“) (15,269,672 unique entries) was performed using Mascot (Matrix Science; version 2.7.0.1), assuming the digestion enzyme trypsin and applying the following parameters: fragment ion mass tolerance 0.5 Da and parent ion tolerance of 10 ppm, up to two missed cleavages as well as methionine oxidation as a variable modification. Scaffold v5.1.2 (Proteome Software Inc.) was used to merge the search results and to validate MS/MS based peptide and protein identifications. During creation of the Scaffold file, an additional X! Tandem search with default settings was performed for validation. Protein and peptide-level false discovery rate (FDR) were set at 5 % and protein groups required to contain at least 1 unique peptide. Protein groups consisted of proteins which were not distinguishable based on identified peptides. For proteomic comparison across samples, the spectral counts were normalized by the protein length, followed by total sample spectral count, as described previously [41].

### Community profiling based on rpS3 marker gene

Genes encoding for the *rpS3* marker gene were identified via hmmsearch [42] against the Phylosift HMM model [43]; PhyloSift Marker: DNGNGWU00028) at an E-value of 1e-28 as well as by pulling out genes annotated as *rpS3* during data processing (see above). All identified *rpS3* sequences throughout all assemblies were then pooled and clustered at 99% similarity as described in Sharon et al. [44] using the usearch -uclust module [45]. Mapping on just the *rpS3* gene would result in inadequate abundance estimates, as reads mapping to the edges of sequences map poorly, causing coverage to drop off at the edges. Hence, for each *rpS3* gene, the scaffold area around the gene was taken into account by extending the scaffold sequences around identified rpS3 genes by up to 1000 bp to either side [46]. The workflow and scripts to identify, cluster, and extract rpS3 sequences as well as calculate their coverage and breadth are provided at: https://github.com/ProbstLab/publication-associated_scripts/tree/main/Figueroa-Gonzalez_Bornemann_etal. This yielded scaffold fragments of about 3500 bp in length, which are expected to result in more accurate abundance estimates than just mapping on the gene by itself. Representative *rpS3* sequences were selected from each *rpS3* cluster by order of the following priorities: 1) centroid and can be extended by 1000 bp into both sides of the gene, 2) not centroid and can be extended into both sides by 1000 bp, and 3) longest scaffold region. The resulting *rpS3* sequences, ideally extended in both directions by 1000 bp, are henceforth called rpS3extended. The taxonomy of the rpS3extended sequences was determined by using usearch (ublast) against the pooled archaeal and bacterial *rpS3* sequences of the GTDB (E-value 0.00001). The best hits from the search were taken, and those rpS3extended sequences without a hit at this threshold were assigned as “Unclassified”. The abundance of the sequences in each sample was determined by mapping of reads back to the rpS3extended sequences and their average coverage per sample was determined. Additionally, the breadth, *i.e.,* the number of nucleotide positions having a coverage of at least 1, was determined. The coverage of an rpS3extended sequence in a given sample was set to zero if the breadth of the sequence in that sample was less than 95%. This was done to ensure that the respective sequences were present in their entirety in the respective sample, instead of just fragments of them. After breadth correction, coverages per sample were normalized by sequencing depth.

### Reconstruction of metagenome-assembled genomes

Reads of all metagenomes were mapped against each assembly to yield differential coverage data used in binning. Metagenome-assembled genomes (MAGs) were reconstructed using ABAWACA v1.0.0 (https://github.com/CK7/abawaca), CONCOCT v1.1.0 (https://github.com/BinPro/CONCOCT), and MaxBin 2.0 v2.2.7 [47]. The bin sets were consolidated with DAS tool v1.1.2 [48] with default parameters. The consolidated bin sets were curated in uBin v0.9.20 [40] and all curated MAGs of all samples were then pooled and dereplicated with dRep[49] at 99% gANI to yield all unique strains present in the dataset. CheckM1 [50] was used to assess and filter genomes based on completeness and contamination, using 70% and 10% as the cutoffs, respectively.

### Taxonomic classification of MAGs

GTDB-tk v.2.1 [51] with the classify_wf workflow against GTDBr207 was used to determine the taxonomic origin of the recovered MAGs. Genes on these genomes were predicted with prodigal in normal mode, either with default genetic codes (for non-Gracilibacteria) or codon code 25 (for Gracilibacteria).

### Generation of protein database from metagenomes

Open reading frames (ORFs) in amino acid format from all metagenomic assemblies, predicted by prodigal in meta-mode, were pooled. ORFs located on scaffolds that had been assigned to Gracilibacteria bins were exchanged with ORFs predicted on the entire Gracilibacteria genome and with codon code 25. Then, all ORFs were uniquified using the *usearch* command [45] to form the database. Each database entry was renamed to Protein_XXXXXXX, with X representing a sequential, unique number for each entry (this database is supplied as Supplementary File S1). Additional metadata, such as whether the respective protein or any of its cluster members were binned and if so, to which bin they belonged as well as their functional or taxonomic annotations, were stored in separate files (see Supplementary File S2).

### Tracking abundances of rpS3 genes and MAGs across samples

Each metagenome was mapped to the dereplicated MAGs or rpS3extended sequences to estimate their abundances across time and filter fractions. Additionally, the breadth of each MAG and rpS3extended sequence in each sample was determined. The breadth is herein defined as the percentage of the sequence length having a coverage of at least 1. This was used to filter out MAGs and rpS3extended sequences that only had coverage because a small proportion of their sequence was shared with other organisms. For genomes, the minimum required breadth to have a valid coverage was set to 30%, for rpS3extended sequences to 95%. If the breadth of a sequence in a sample was below this threshold, the coverage was set to zero.

### Normalization of abundances of MAGs and rpS3 genes

MAG and rpS3extended abundances were normalized to the sequencing depth per sample using the following formula:

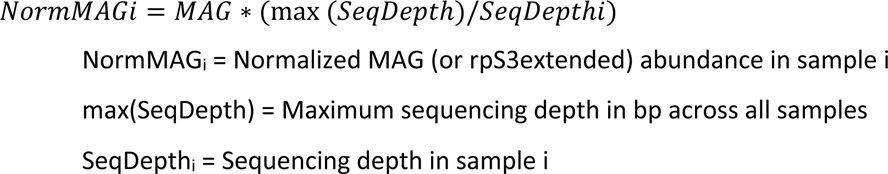

### Shannon and Simpson diversity estimates of assemblies

Shannon and Simpson diversity estimates were calculated from rpS3extended sequence abundance tables using the diversity function of the vegan R package (https://github.com/vegandevs/vegan).

### Rank abundance estimates of phyla

The normalised coverage of rpS3extended sequences assigned to the same phylum based on GTDB annotation were summed up by sampled time-point and by each filter fraction (0.1 µm or 0.2 µm).

### Non-Metric Multidimensional Scaling (NMDS)

NMDS, based on Bray-Curtis dissimilarities, were calculated using the *metaMDS* function of the *vegan* package, using species tables of either MAGs or rps3extended sequences. All samples were analyzed while being pooled together as one database, as well as split into the two filter fractions. Only NMDS plots with a convergent solution (after a maximum of 400 tries) as well as sufficiently low stress are shown.

### Proportionality networks

The temporal dynamics of MAGs across time in the two filter fractions were investigated using proportionality networks implemented in the R package *propr* [52]. In contrast to conventional correlation analyses, proportionality networks are not biased when relative data is compared [53]. The proportionality measure rho (ρ) was employed and cutoff thresholds were determined to correspond to False-Discovery-Rate (FDR) thresholds of 1% and 5%, respectively, using the *updateCutoffs* function.

### Metabolic analyses

Metabolic potential of the microbial community was determined using METABOLIC [54]. Genome annotation of Gracilibacteria was done using the Genoscope platform MAGE [55]. In-depth metabolic potential was determined by combining the annotation information from MAGE with information from MetaCyc [56], KEGG [57], and UniProt [58] (see Table S7 for automatic annotation results). Furthermore, unclassified proteins were manually searched against protein domain databases of NCBI protein domains (https://www.ncbi.nlm.nih.gov/Structure/cdd/cdd.shtml) and AlphaFold-EMBL database [(https://www.ebi.ac.uk/Tools/sss/fasta/), see Table S8 for manual annotation results). The information obtained in the analyses of automatic annotation was compared with the metaproteomics data to evidence expressed proteins.

### Comparison of available microbial populations from cold-water geysers

All dereplicated MAGs from Crystal Geyser (Utah, USA) [59], Geyser Andernach (Andernach, Germany) [60] and the Geyser Wallender Born (Wallenborn, Germany) were pooled and compared via FastANI[61] with a >= 50% coverage.

### Visualization

Figure panels, including the NMDS, were generated with *ggplot2* [62] in R [63], and subsequently assembled and post-processed in AffinityDesigner V2 (https://affinity.serif.com/). The phylogenetic trees produced by GTDB-tk were visualized using Dendroscope v3.5.10 [64] and processed with iTol v6 [65]. The proportionality networks were generated using the *iGraph* package [66]. The number of shared genomes between the cold-water geysers were analyzed using *ggupset (*https://github.com/const-ae/ggupset). The metabolic reconstruction scheme was generated manually in Affinity Designer V2.

## Results and discussion

### Microbial community of the geyser is dominated by Gallionella and Gracilibacteria

Size fractionation of 22 groundwater samples discharged from Wallender Born resulted in 15 samples on 0.2-µm filters, 14 samples on 0.1-µm filters, and one bulk 0.1-µm filter sample. The subsequent metagenomic sequencing resulted in 725 Tbps of Illumina NextSeq data.

These data covered 78% of the microbial diversity (on average) in the samples based on Nonpareil3. After individually assembling the metagenomes, we determined that about 80.0% (+-3.2% STDEV) of the reads mapped back to the scaffolds, with only slight differences between 0.1-µm and 0.2-µm filter fractions (78.2% +-2.9% STDEV and 82.6% +-1.9% STDEV, respectively; see Supplementary Table S4 for reads, assembly and Nonpareil3 statistics). We hence conclude that the assemblies are representative of the microbial community of the geyser. The Shannon diversity index based on *rpS3* marker genes was on average 3.22 for 0.1-µm size fractions, and 3.50 for 0.2-µm size fractions, while the Simpson diversity index was on average 0.90 for both size fractions with similar intra-sample variation (0.42 to 0.11, and 0.30 to 0.01, respectively; Figure 1A). Based on *rpS3* abundances (Figure 1B), the communities were dominated by Proteobacteria (58.0%) followed by Patescibacteria (26.2%) and Bacteroidota (4.57%; please refer to Supplementary Figure S3 for the complete rank abundances and Supplementary Figure S4 for phylum-level taxa abundances across time-series samples). NMDS analyses revealed that the 0.1-µm filter fractions showed a great intra-group dissimilarity, and that 0.2-µm samples and the bulk sample clustering in close proximity (Figure 1C). Individual NMDS analyses of 0.1-µm and 0.2-µm filter fractions confirmed this finding, with the NMDS of the 0.1-µm samples being much more dissimilar compared to their 0.2-µm counterparts (Figure 1C).

**Figure 1.**
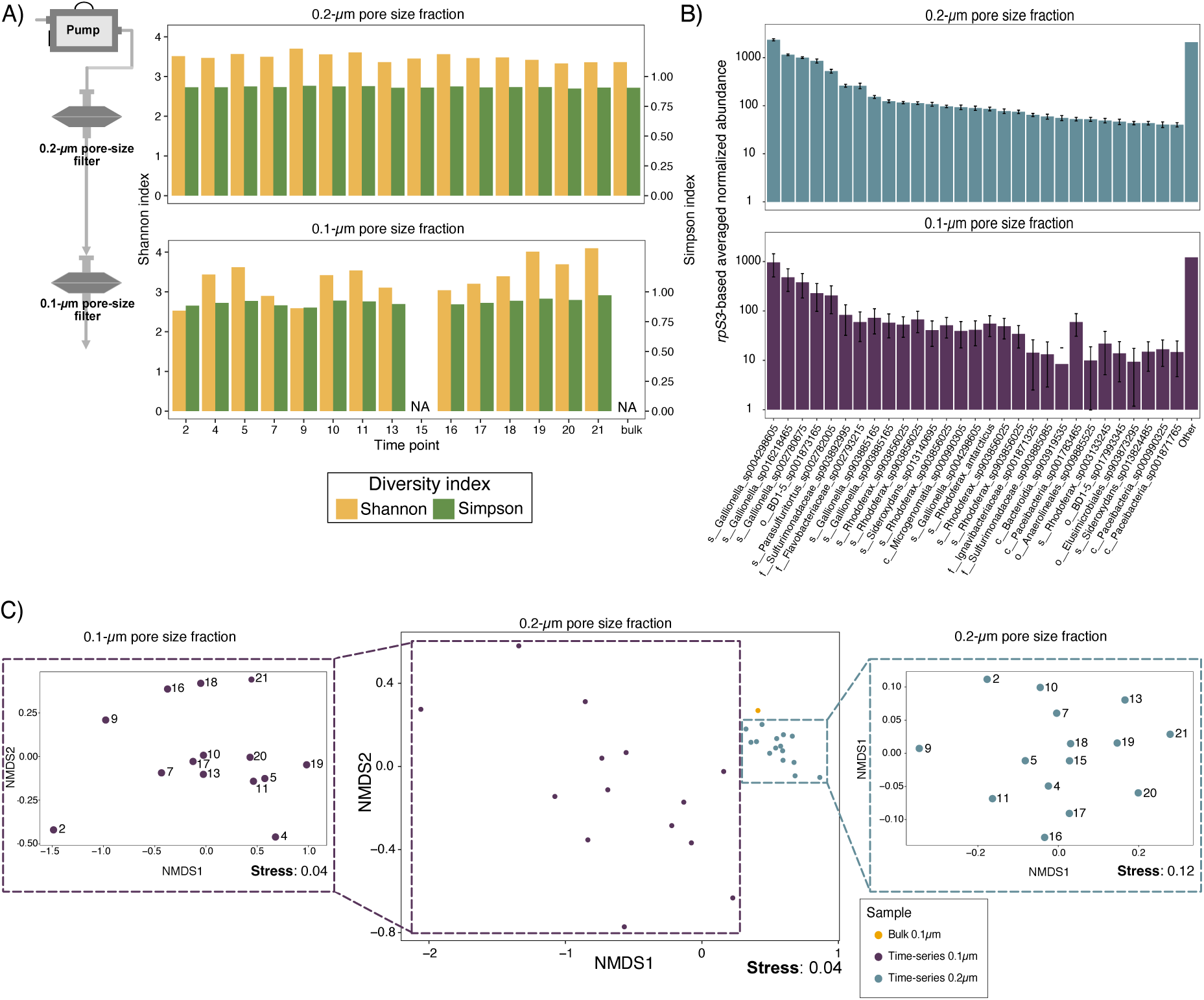
Microbial community composition of the geyser Wallender Born. **A)** The scheme to the left side illustrates the principle of sequential filtration we used for this study. The bargraph on the right displays the Shannon and Simpson diversity indices for the 0.2-µm and the 0.1-µm filtered fractions. NA denotes samples that we were unable to sequence due to low biomass. **B)** Normalized average abundance, based on *rpS3* gene, for both the 0.2-µm and the 0.1-µm filtered fractions. The taxonomy is based on GTDB-tk. The figure illustrates the top 29 species with the rest of the species pooled under “Other”. Error bars denote standard deviation (not shown for all low abundant organisms grouped as Other). **C).** NMDS, based on *rpS3*, of both filtered fractions, and individual NMDS for the 0.2-µm and the 0.1-µm fractions. Central panel depicts the NMDS of both filters, while side panels show separate NMDS for 0.1-µm and 0.2-µm filters, respectively.

### Metaproteogenomics reveals potential key stone organisms in the geyser community

We reconstructed 751 metagenome-assembled genomes (MAGs), with completeness of at least 70% and contamination of at most 10%, based on CheckM1 estimates. These MAGs belonged to 123 strain clusters after dereplication at 99% ANI (see Supplementary Table S6 for genome information). Phylogenetic analysis based on GTDB-tk indicated that these 123 dereplicated MAGs belonged to 26 different phyla, including Nitrosomonadales, Proteobacteria, Nitrospirae and Bacteroidetes, and multiple bacteria of the Candidate Phyla Radiation (CPR), in particular Nomurabacteria and Gracilibacteria (Figure 2). Microbial communities, in particular those of the 0.2-µm filter fraction, were stable across the sampled time series, with the filter fraction being the main discriminatory factor for microbial communities (Figure 3A). Ordination analyses based on relative abundance of MAGs agreed with *rpS3*-based ordinations, displaying that there was a significant difference between the communities on the 0.1-µm filters compared to those of the 0.2-µm filters, which were comparatively stable (Figure 1C). Comparison with known cold-water geyser microbial communities [59,60] (Geyser Andernach (GA), Germany; and Crystal Geyser (CG), USA) showed that most community members were specific to their respective ecosystems, with only Altiarchaea and Hydrogenophilales being present in all three ecosystems (Supplementary Figure S5). While Altiarchaea were the dominant organisms in both GA and one aquifer in CG, they constituted a minority in WB. Out of the 123 WB genomes, 24 had related populations based on FastANI (>= 50% genome coverage, >= 75% ANI, [61]) in the CG ecosystem while only a single population genome was related between the WB and GA populations, which might relate to the larger diversity of the CG ecosystem compared to the other two ecosystems.

**Figure 2.**
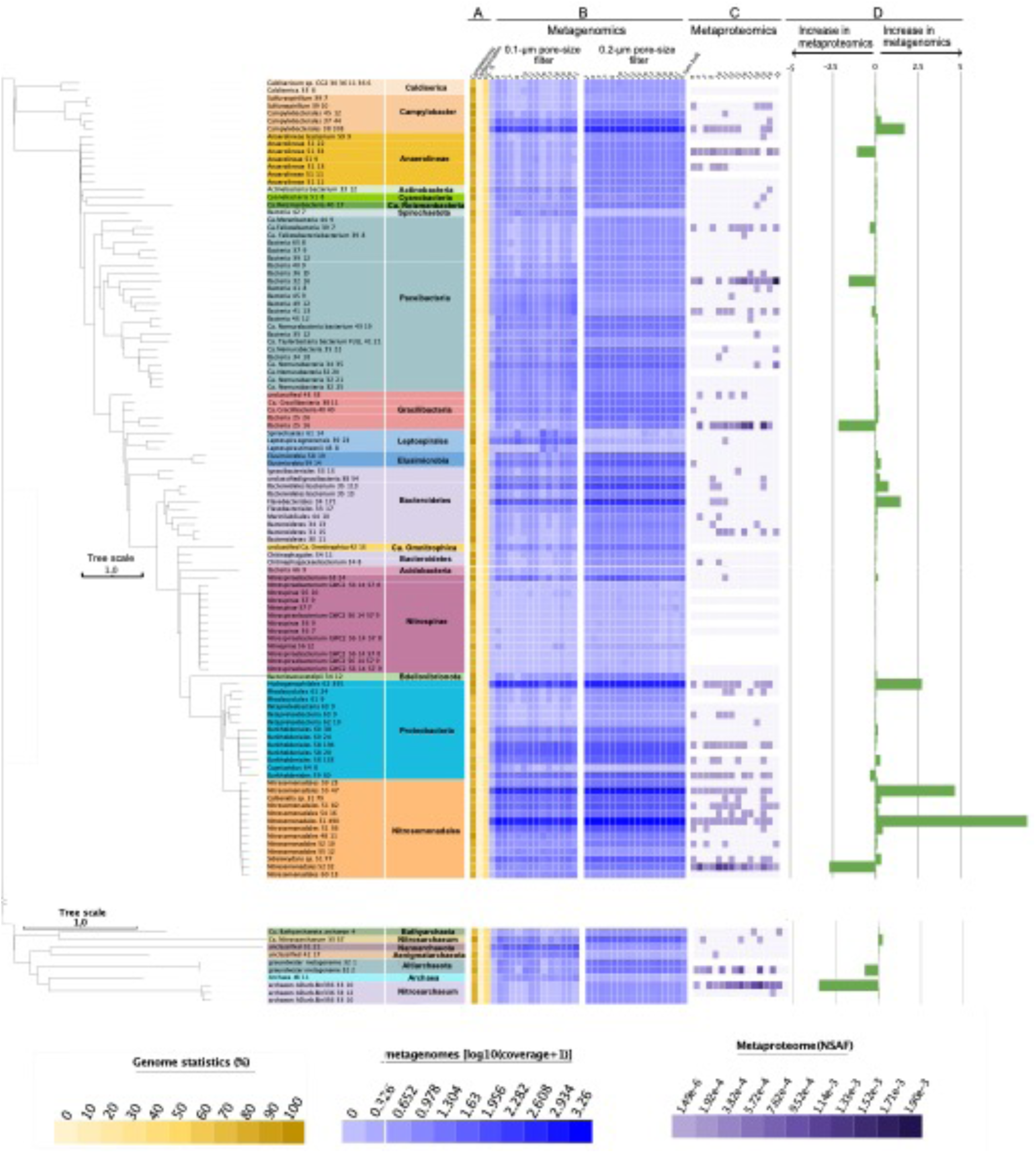
Phylogeny of high-quality genomes and their prevalence in metaproteogenomics. Colored sections to the right of the phylogenetic tree denote the phylum-level taxonomic classifications of high-quality genomes. **A)** Yellow-colored heatmap indicates the genome statistics (completeness, contamination, and GC % content). **B)** The blue heatmap shows the abundance of the genomes throughout the time-series, separated into 0.1-µm filter and the 0.2-µm filter fractions. **C)** The purple heatmap conveys the abundance of the genomes based on the summed counts (NSAF) of proteins belonging to the respective genomes. **D)** Green bar charts denote the difference in the median relative abundance of the metagenomic-based genome abundance relative to the proteomic-based genome abundance on the 0.2-µm filters. The median relative abundance was calculated by setting the total sample abundance to 100%, both in metagenomics as well as in metaproteomics, and afterwards calculating the median percentage per genome. Negative numbers indicate higher relative abundance in metagenomic data than in the metaproteomic dataset, and vice versa for the positive percentage numbers.

**Figure 3.**
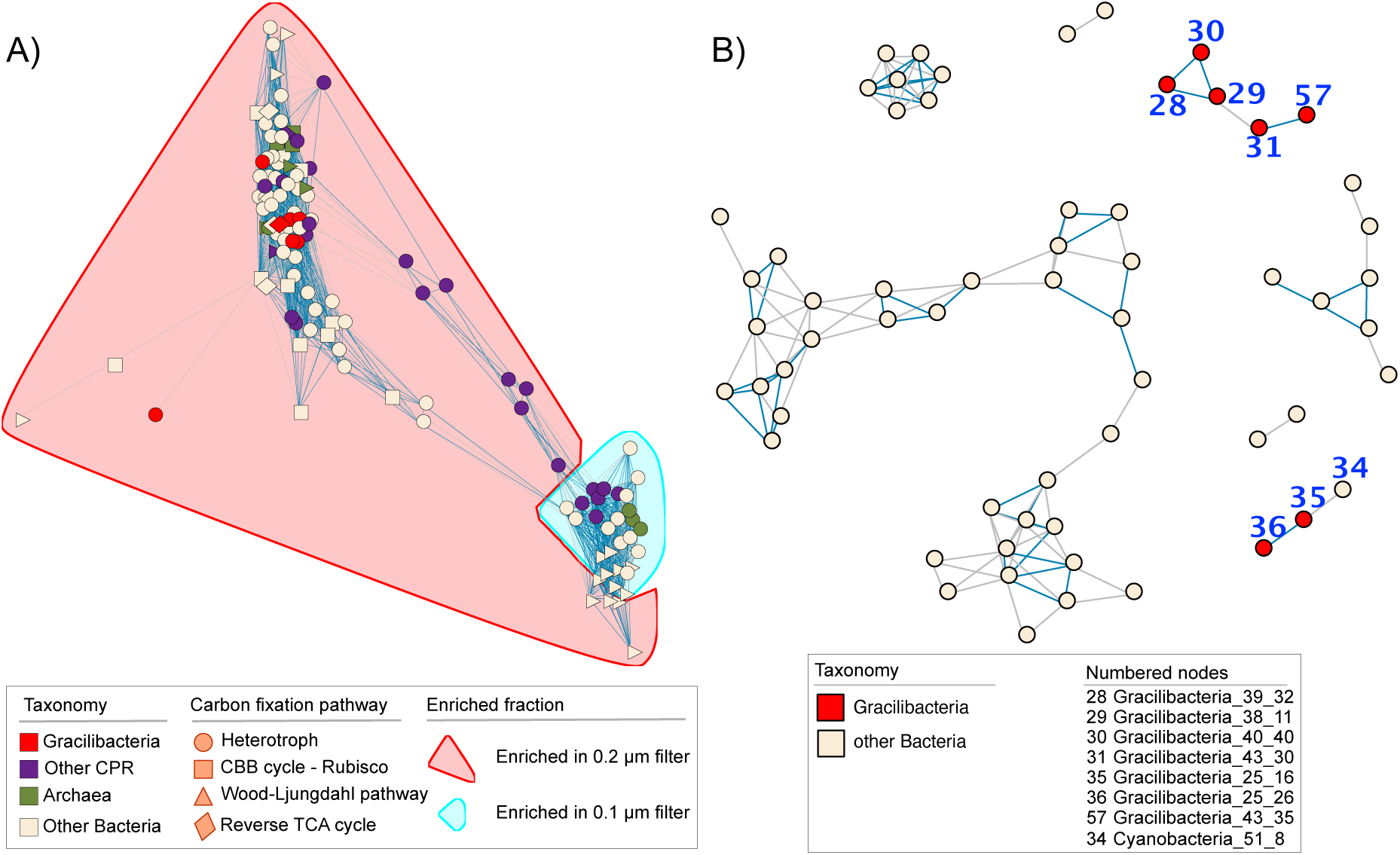
Co-occurrence networks of unique genomes. **A)** Co-occurrence of the overall prokaryotic community of WB; color of shape denotes taxonomy, shape relates to carbon fixation pathway detected in each genome, enrichment of genomes in either the 0.2-µm or 0.1-µm filter fraction is shown by the areas highlighted in red or blue. **B)** Co-occurrence network of 0.2-µm filter samples focusing only on Gracilibacteria and non-CPR bacteria. Blue connecting lines indicate 1% FDR, while grey lines signify 5% FDR. Gracilibacteria were co-correlated and showed only one weak correlation with a cyanobacterial MAG.

Although other high-CO_2_ geysers have been metagenomically analyzed in the past [16,60,67], little is known about the activity or the contribution of the organisms to the protein pool of the community. To elucidate the expression profile (*i.e.*, expressed phenotype and metabolic functions) of the detected organisms in the metagenomes of geyser WB, we performed metaproteomics of the very same samples that were analyzed for metagenomics (see Supplementary Figure S2 for sample distribution of metagenomics and metaproteomics). Metaproteomic datasets showed a batch effect in the NMDS analysis, with the three categories of samples (time-series, size, and bulk) clustering separately in the ordination (Supplementary Figure S6). Subsequently, we focused our analyses on the 0.2-µm fraction only and compared metaproteomic and metagenomic data of these samples. Across time, the proteomics profile of the community in the 0.2-µm fraction was more dynamic than the relatively stable genome abundances, with only few organisms showing consistent expression throughout the time series (Figure 2B-D). Only the most abundant organisms in metagenomics (including a Campylobacter, a Hydrogenophilales, and two Nitrosomonadales closely related to Gallionella) were also fairly consistently represented in the proteomic datasets (Figure 2). However, the most abundant members based on proteomics included organisms of the Anaerolineae, a Paceibacteria, a Gracilibacteria, as well as some Nitrosomonadales, which were all only minor community members in the metagenomic dataset (Figure 2B-D). Based on their consistently high abundance in both metagenomes and metaproteomes, we conclude that these CPR bacteria likely represent key organisms in the microbial community of the geyser.

### Gracilibacteria are enriched in the 0.2-µm fraction and show no co-correlation with other prokaryotes

CPR bacteria, in particular Nomurabacteria and Gracilibacteria, represented 31 of the 123 unique MAGs recovered from geyser WB. Due to their predicted symbiotic lifestyle [4,8], we sought to reconstruct co-occurrence networks to establish potential symbiont-host relationships. As correlating relative abundance data, such as genome abundances, is problematic due to relative data by default fostering significant correlations [53], we utilized the proposed alternative proportionality [53], implemented in the R package ‘*propr’* [52]. The resulting network showed two clusters of organisms (Figure 3A), which has previously been suggested to be related to different aquifers and thus groundwater sources at Crystal Geyser [59,68]. Nonetheless, further analyses showed that the clusters observed for WB are due to the fractionation, where microbes group based on their enrichment pattern in either the 0.1- µm or 0.2-µm filter fraction. We observed that the WB Gracilibacteria genomes are enriched in the 0.2-µm fraction, while other CPRs were either distributed between both network clusters or situated between the clusters.

When focusing the analysis on the dataset from 0.2-µm filters, on which both hosts and host-symbiont attachments should accumulate, the proportionality network showed only one weak link for one out of seven Gracilibacteria with another organism, a Cyanobacterium (genome Cyanobacteria_51_8; 5% FDR; Figure 3B). *Vampirococcus*, closely related to the WB Gracilibacteria (Figure 4A), have been reported to predate on autotrophs such as *Chromatium* and *Halochromatium* [23,69]. Although there is one report of viable Cyanobacteria surviving on a hydrogen-based metabolism in a 613-m-deep borehole in Spain [70], these organisms can be contaminants of open geysers [60]. While these Cyanobacteria may still fall prey to Gracilibacteria, correlation and proportionality networks have previously been discussed to be biased towards organisms with scavenging lifestyles (based on Lotka-Volterra models; [71]). This reported bias, and given the absence of co-correlation with Gracilibacteria, we suggest that the detected Gracilibacteria populations follow a parasitic lifestyle as previously described for the closely related Abscondidabacteria [23,24].

**Figure 4.**
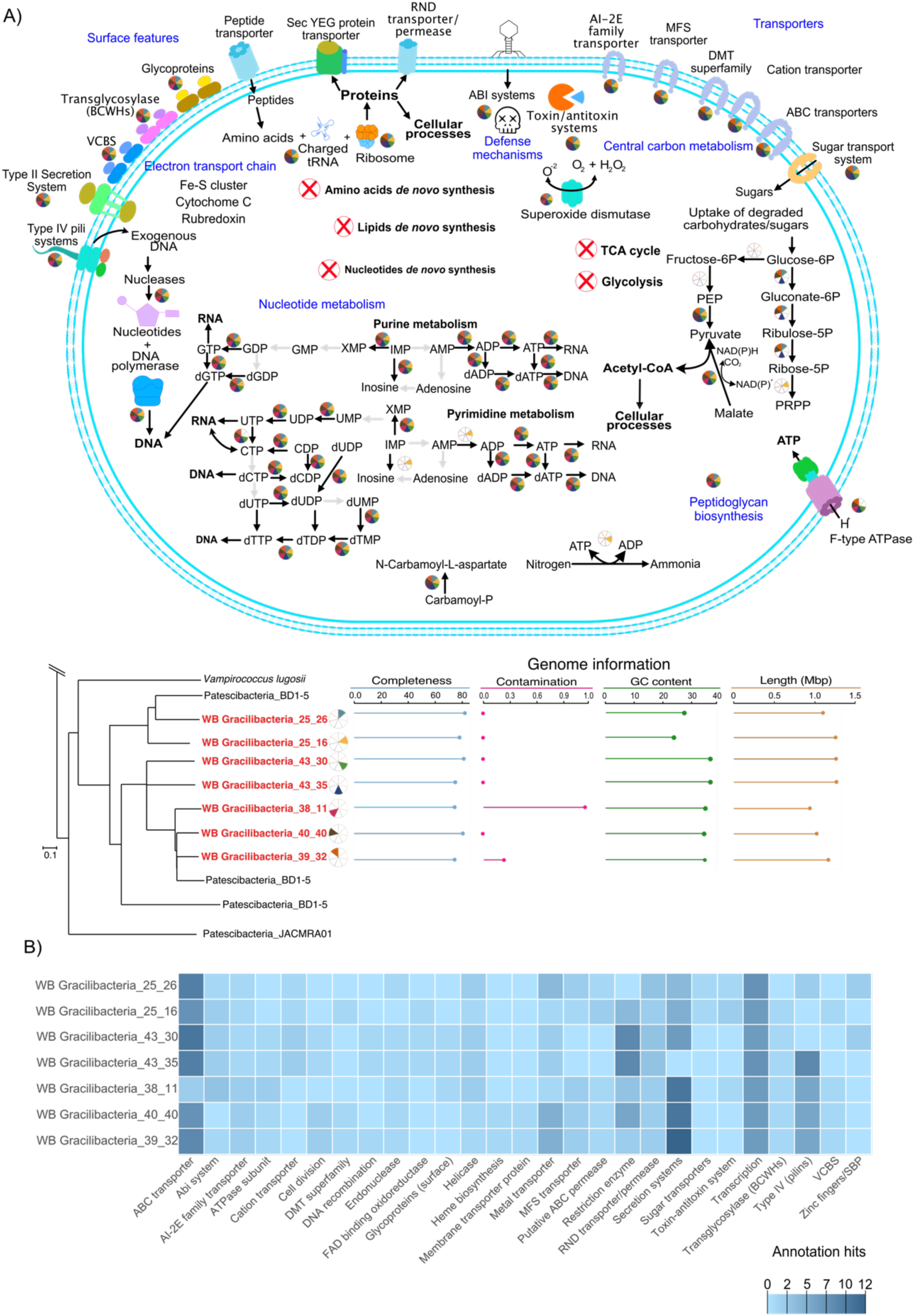
Metabolic overview of the seven WB Gracilibacteria genomes. **A)** Schematic metabolic overview of the major traits found in the WB Gracilibacteria and phylogenetic placement of the seven genomes along with genome statistics. Please note that the individual wedges of the pie-charts relate to the presence of individual metabolic steps encoded in the different genomes, depicted in the metabolic scheme. **B)** Specific proteins of interest that were identified in the genome annotation of Gracilibacteria.

### Metaproteomics suggests an active symbiotic/predatory lifestyle of WB Gracilibacteria

WB Gracilibacteria missed all of the steps related to glycolysis, with the exception of genes encoding for the conversion of phosphoenolpyruvate (PEP) to pyruvate (Figure 4A, Supplementary Figure S7). The latter could enable the generation of ATP via substrate level phosphorylation using pyruvate kinase (EC. 2.7.1.40), as previously hypothesized for *Vampirococcus lugosii*, a cultivated member of the related Abscondidabacteria [23]. Additionally, WB Gracilibacteria MAGs encoded for a variety of electron carrier proteins (*e.g.*, Cytochrome C, Fe-S cluster proteins, rubredoxin, peroxidases) and F-type ATPase subunits. Given the presence of electron transport systems, the WB Gracilibacteria could establish a proton gradient to make use of the ATPase for energy generation and use the electron carrier proteins to maintain oxidative stress under control [23]. Energy generation via the use of ATPase is further evidenced by the F-type subunits identified in the proteomic analyses (see Supplementary Figure S8, “ATPase” category). By contrast, it has been argued that CPR bacteria in general could take electrons from their host and use ATPase in a reverse manner (consuming ATP) to drive their antiporters [23], aiding in the uptake of external molecules. Given the patchiness of their central carbon metabolism, the annotation of two enzymes in the TCA cycle, namely pyruvate formate lyase (activating enzyme, EC. 1.97.1.4) and ATP citrate (pro-S)-lyase (EC. 2.3.3.8), suggests that WB Gracilibacteria need to fill metabolic gaps with externally derived compounds, such as citrate.

The seven Gracilibacteria genomes also encoded for interconversion steps for the purine and pyrimidine biosynthesis pathways, with one genome (WB Gracilibacteria 26_16) encoding for the conversion of ribose-5P to phosphoribosyl diphosphate (PRPP) (Figure 4A). The pyrimidine interconversion steps start with UMP/dUMP and ultimately yield CTP/dCTP that can be used to synthesize RNA and DNA, respectively. As for purines, the conversion steps go from IMP to ATP and dATP which, in a similar manner as pyrimidines, can serve as substrate for the synthesis of RNA and DNA, respectively. While PRPP is an important metabolic intermediate, in particular for the biosynthesis of nucleotides [72], most of the WB Gracilibacteria do not encode for its synthesis (Figure 4) suggesting that this intermediate has to be retrieved from external sources.

The limited metabolism of the WB Gracilibacteria suggests that these bacteria require molecular building blocks from other organisms (refer to Supplementary Figure S7 and Table S7 for the full KEGG annotation of these genomes), but it remains unclear how they obtain these necessary biomolecules. Given that no potential host(s) for the WB Gracilibacteria could be identified using co-occurrence patterns, as well as their extremely limited central metabolism, we set out to investigate their potential lifestyle based on metaproteogenomics, focusing on (trans)membrane and cell surface features (see Figure 4B and Supplementary Table S8 for proteome annotations of Gracilibacteria MAGs). We identified membrane-associated proteins predicted to be functioning as transporters, in particular ABC transporters, which aid in the uptake of a broad spectrum of molecules from the surrounding of the cell [73–76]. Other transporters identified in the genome, such as the DMT superfamily and major facilitator superfamily (MFS), enable the transport of drugs and a broad range of molecules across the membrane, including antibiotics [77]. Consistent with other Gracilibacteria, the WB Gracilibacteria expressed type IV pili systems,which can aid in the uptake of DNA from the environment [78,79] and have been shown to be essential for host adhesion in the case of TM7i and its *Leucobacter aridicollis* host [80]. The DNA uptake is further supported by the observed strong expression of endonucleases. Consequently, WB Gracilibacteria were predicted to attach to their host or make use of their appendages to obtain molecules, which is likely necessary during the turbulent eruptions of the geyser. Furthermore, most WB Gracilibacteria encoded for the resistance-nodulation-division (RND) family of proteins, which also have transporter functions and efflux activity [81] but were not detected in the proteome. Two genomes (WB Gracilibacteria 25_16 and 25_26) encoded for a putative Vibrio, Colwellia, Bradyrhizobium, and Shewanella (VCBS) repeat-containing protein, a group of proteins that are proposed to contain domains with functions related to cell adhesion (InterPro entry: IPR010221). We were able to identify them in the proteomic data by means of manual search of the unclassified proteins (Supplementary Figure S8, “Adhesion” category). Translocases and peptidases identified in the proteome could aid in obtaining amino acids probably to mainly carry out protein synthesis but also other reactions, like deamination, and mobilization of molecules across the membrane. Strikingly, we identified a vast number of type II and IV secretion systems in all seven genomes, of which some subunits were also identified in the proteome (Supplementary Figure S8, “Secretion system” category). Type IV secretion systems are related to pili and its assembly subunits [82–84], while secretion systems II are used by Gram-negative bacteria to excrete proteins and other molecules that aid in signaling or microbial pathogenicity [82,85–88]. Further evidence for the active expression of type II secretion systems in Gracilibacteria is their differential expression when comparing the most abundant proteins between the 0.2-µm filter and the 0.1-µm filter, showing that subunits for these secretion systems are highly expressed by WB Gracilibacteria (Supplementary Figure S9). We also identified, in both genome and proteome data, subunits of the translocase Sec system, which is important for the maturation and secretion of proteins and their insertion into the inner membrane [82,89–91]. Two proteins were annotated as transglycosylase, which are involved in cell wall interactions, such as anchoring and even cell lysis [92–94]. Added to these cell surface features, the WB Gracilibacteria also encoded for a complete peptidoglycan biosynthesis.

Publicly available Gracilibacteria genomes are missing steps in central metabolic pathways, such as glycolysis and the tricarboxylic acid cycle (TCA), but encode for the complete oxidative branch of the pentose pathway [23,24,28]. Likewise, the WB Gracilibacteria showed similar traits and encode for several cell surface features associated with a predatory lifestyle (Figure 4). Annotations included gene functions related to transporters (*e.g*, ABC, protein exporters, efflux systems), secretion systems type II and IV, pilus and flagella assembly, oxidoreductases, peptidases, DNA repair, peptidoglycan assembly, and nucleotide interconversions, all of which agree to the previously described metabolic patterns encoded in cultivated and uncultivated Gracilibacteria [23,24].

Taken together, the presence of the abovementioned findings suggests an active predatory lifestyle in the groundwater fluids discharged by the Geyser, where the WB Gracilibacteria obtained their needed biomolecules by means of adhesion using pili and flagella, subsequently lysed the target cell using their surface features such as the injection and efflux systems, and ultimately made use of their transporters and appendages repertoire to take in the cellular components released. Once taken in, the range of enzymes (*e.g.,* endonucleases, proteases, metalloenzymes) in the Gracilibacteria would be in charge of breaking down big molecules and incorporating smaller molecules into its metabolism.

### Gracilibacteria as key members of the microbial community of geyser Wallender Born

While the functional significance of Gracilibacteria in the geyser’s ecosystem is still unclear, their symbiotic lifestyle coupled to their great abundance in metagenomics and metaproteomics demonstrates that they are key players in the geyser’s community. Taking into account that cell counts for the 0.2-µm filter samples ranged between 2.07x10^4^ to 5.94x10^4^ cells ml^-1^ (Table S1), cell numbers that were on the low end for a shallow subsurface environment [95], and considering the regular flushing of the geyser’s ecosystem (eruptions occur in 30-40 minute intervals), it can be assumed that availability of molecules is low. Added to this, it has been proposed that the small cell size of CPR, in this particular case of Gracilibacteria, is an advantage in oligotrophic environments since the increased surface-to-volume ratio of the cell is optimal for the uptake of nutrients [96,97]. Additionally, the small cell size would allow for more Gracilibacteria to target the same host cells [21]. Considering that most of the Gracilibacteria genomes did not co-correlate with other prokaryotes in the community, we assume that the Gracilibacteria can target a broad range of microbes to feed on. This assumption and the high abundance of Gracilibacteria compared to other members of the overall geyser’s microbial community (Figure 2) indicates that the Gracilibacteria thrive in this environment, likely by means of attaching to other microbes to predate, as opposed to scavenging in such a low-nutrient environment. If one considers a similar case as with *Ca.* A. praedator, which is proposed to play a role in population control by a “Kill the Winner” approach [24], Gracilibacteria in WB could potentially target other organisms in the geyser, lysing them and effectively keeping any one species from outcompeting the rest of the microbial community. In oligotrophic environments, such as geyser WB, it is expected that predators, deemed “competition specialists”, dominate the community since competition is a selective pressure [98]. It should be noted that the “winner” in such scenarios is not necessarily the most abundant organism (*e.g.,* Gallionella in WB) but rather the competition specialist [98], in this case the predatory Gracilibacteria or key stone organisms that are low abundant (infrequent targets) but highly active. We propose that Gracilibacteria likely serve as a population control in WB, aiding in maintaining the dynamics of the geyser’s microbiome stable throughout time.

## Conclusions

We present an in-depth metaproteogenomics study of a groundwater microbial community accessed through a cold-water geyser. Metagenomic abundances show that the microbial community is dominated by Nitrosomonadales, specifically by members of Gallionella spp., Campylobacter and Hydrogenophilales, which also showed expression in the metaproteomic data. Nonetheless, metaproteomics showed that Anaerolineae, Paceibacteria, Nitrosomonadales, and in particular Gracilibacteria were the most active in terms of protein abundance. Analyzing metaproteogenomic data, we were able to draw conclusions regarding the lifestyle of Gracilibacteria. They show metabolic overlaps with other members of the Gracilibacteria lineage, with very limited central metabolism, while encoding for interconversion steps of nucleotide biosynthesis and peptidoglycan synthesis. However, their core metabolism patchiness is compensated by encoding and expression of several types of transporters, membrane-related proteins, and enzymes. Added to this, we identified proteins that point towards cell-cell interactions and type II and type IV secretion systems. Based on these results, we propose that the WB Gracilibacteria follow a predatory lifestyle, likely being capable of lysing other members of the geyser’s microbial community. Considering the membrane features we identified and the lack of identification of a potential host using co-occurrence patterns, we hypothesize that the WB Gracilibacteria are predators that rely on other members of the community to obtain molecular building blocks to survive. At the same time, WB Gracilibacteria potentially assist in population control for the microbial community of WB geyser. Additionally, they have the potential to become a liability to the community if nutrients deplete too much or too fast, or if stress conditions become challenging, during which their predatory lifestyle would be an advantage to survive. The results of this metaproteogenomic study focusing on groundwater Gracilibacteria provides *in situ* evidence regarding the activity of Gracilibacteria suggesting a predatory lifestyle in oligotrophic groundwater ecosystems.

## Supporting information

File S3: GTDB-tk classification of bacterial MAGs

File S4: GTDB-tk classification of archaeal MAGs

## Declarations

### Ethics approval and consent to participate

Not applicable

### Consent for publication

Not applicable

## Acknowledgements

We would like to thank Ines Pothmann for laboratory maintenance, Ken Dreger for server administration and maintenance, as well as Rashi Halder of the Luxembourg Centre for Systems Biomedicine (LCSB) for sequencing. PAFG and TB would like to thank Diana and Frank for their hospitality during the sampling campaign, as well as the Wallender Born municipality for access to sampling the geyser. We would also like to thank the Aquatic Microbiology group for allowing us the use of their ion chromatography instruments. Sebastian Grund is acknowledged for technical assistance in proteomics sample preparation and Anke Trautwein-Schult for support in metaproteomics data processing.

## Availability of data and material

Raw sequencing data and MAGs used in this study from geyser Wallender Born have been deposited at SRA and Genbank, respectively, and are available under the BioProject PRJNA1001268. The mass spectrometry proteomics data have been deposited to the ProteomeXchange Consortium via the PRIDE [99] partner repository with the dataset identifier and the password goTjmuGe.

## Author contributions

P.A.F.G. and T.L.V.B. carried out sampling, DNA extractions, and cell counting. T.L.V.B. performed geochemical measurements, assisted by T.H. during TOC measurements. P.A.F.G. did the metabolic and proteomic analyses, data interpretation and prepared the corresponding figures, as well as preparation of samples for protein extractions. T.L.V.B. performed bioinformatics and statistical analyses, and developed the relevant figures, as well as carried out the geochemical measurements. T.H., S.M., and D.B. conducted proteomic extraction and measurements, as well as data interpretation of it. J.S., C.M., S.P.E., and J.P. aided in the metagenomic analyses and binning. U.S. provided geological information. B.B. aided in discussion and insight into data interpretation. A.J.P. conceptualized the study. P.A.F.G., T.L.V.B. and A.J.P. wrote the manuscript with revisions from all co-authors.

