## Supplementary Information for "Metaproteogenomics resolution of a high-CO_2_ aquifer community suggests an active symbiotic lifestyle of groundwater Gracilibacteria"

Main Supplementary File for:

*authors contributed equally

**Content:**

1. Supplementary Figures
2. Supplementary Tables
3. Supplementary Files

### Supplementary Figures

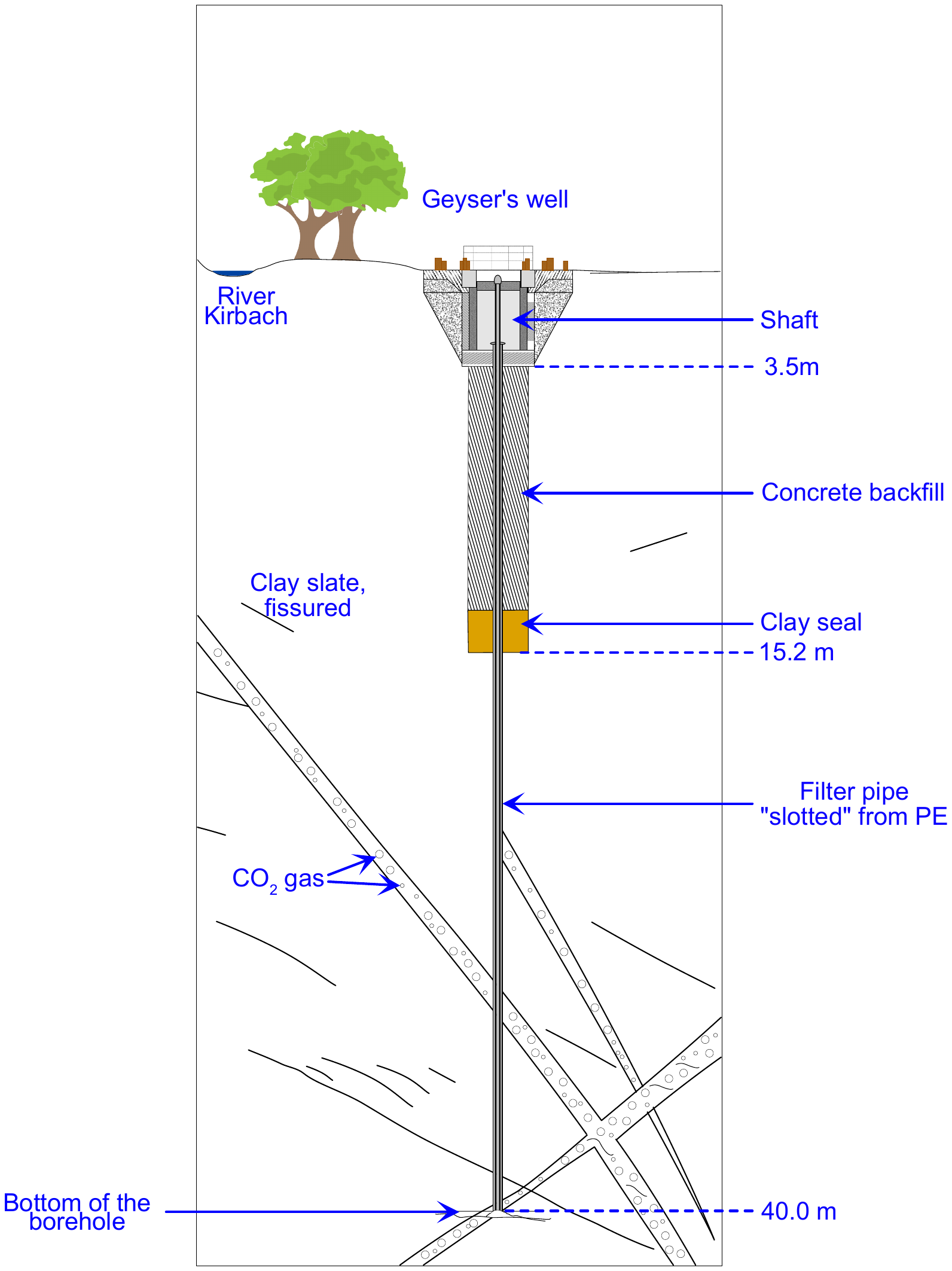

**Supplementary Figure S1. Scheme of the subsurface and borehole of the cold-water geyser Wallender Born.** The borehole has a length of 40 m, with a clay seal at around 15 m followed by a concrete backfill. The geyser has a shaft area of about 3.5 m in depth, where groundwater accumulates after every eruption. The eruptions of the geyser are dependent on the accumulation of CO_2_ in the groundwater. This CO_2_ derives from a magma chamber that lays at great depth in the subsurface (not pictured). The gas rises through cracks and fissures and once it comes into contact with groundwater, the water’s pressure and low temperature allow large amounts of CO_2_ to dissolve in it. When the groundwater is saturated with CO_2_, bubbling occurs, and the pressure of the water column starts to decrease near the surface. This decrease in pressure causes the degasification of the CO_2_ that had dissolved in the groundwater, and gas bubbles find their way to the surface. The gas-water mixture erupts releasing the excess CO_2_. Once the bubbling stops the cycle begins anew, with eruption occurring every 30 to 40 minutes (Translated and adapted from: Information Board on-site *Eifelgemeinde Wallenborn “Der Wallenborn”*).

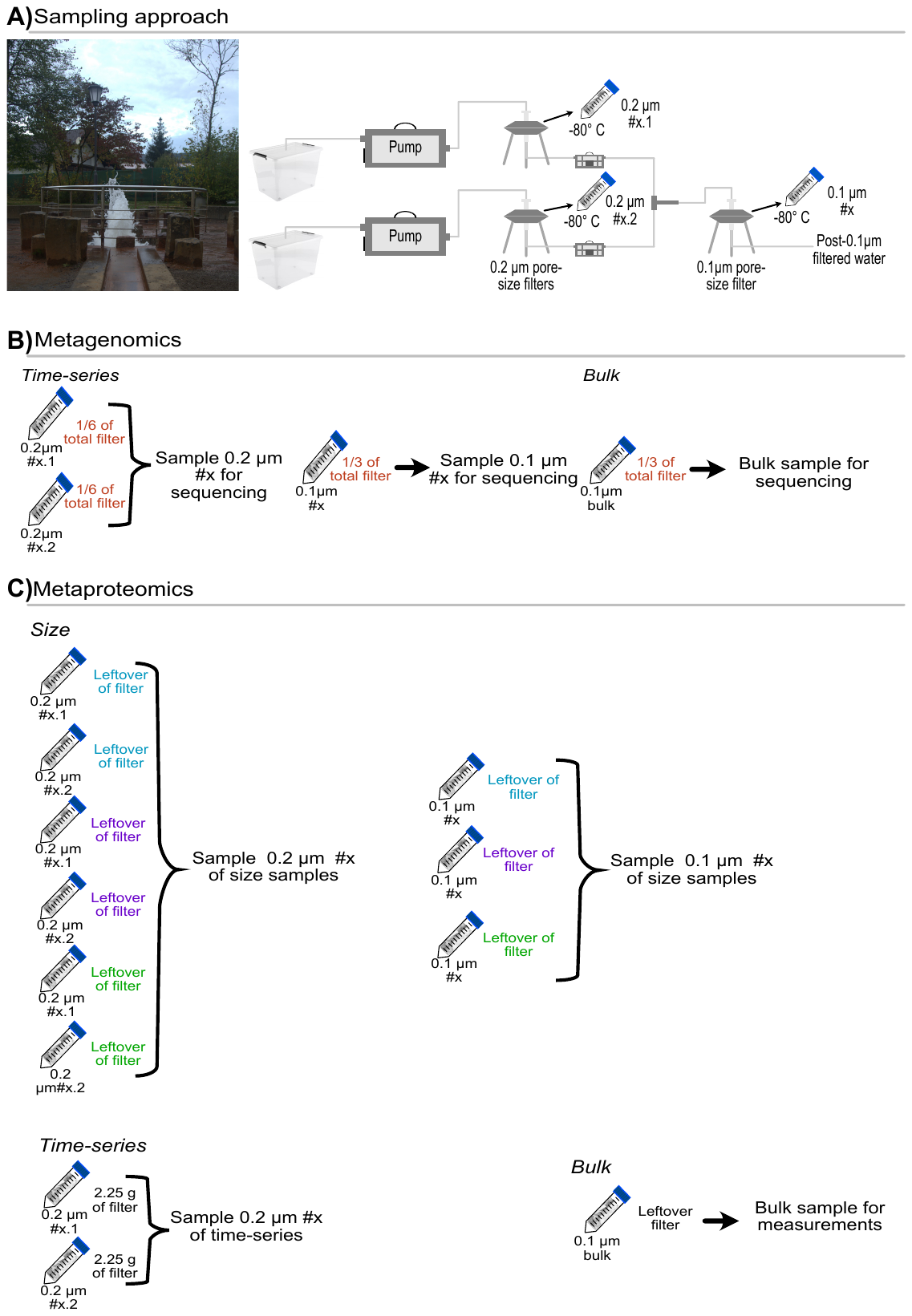

**Supplementary Figure S2. Sampling approach and sample distribution for metagenomic sequencing and metaproteomic measurements. A)** Scheme of the sequential filtration, where groundwater is collected in sterile containers that lead to two pumps, and each pump is in turn connected to a 0.2-µm pore-size filter. The flow-through from both 0.2-µm filters is combined for subsequent filtration through one 0.1-µm pore-size filter. This way, each 0.1-µm filter sample is made up from a pair of 0.2-µm filters samples. Volume counters downstream of both 0.2-µm filters are used to keep track of their respective contribution to the 0.1-µm filter. Roughly 100 L were collected on each 0.2-µm filter per time point (see table S1 for precise volumes). **B)** Sample distribution for metagenomic sequencing of the time-series and bulk samples. For each timepoint of the time-series, 1/6 of the total frozen mass (in g) of each 0.2-µm filter pairs were taken and pooled as one sample for DNA extractions. The corresponding 0.1-µm filters were handled in the same manner, with 1/3 of the frozen mass taken for DNA extraction. The same was done for the bulk sample, taking 1/3 of the 0.1-µm bulk filter for sequencing. **C)** Metaproteomics pooling of the 0.2-µm and 0.1-µm filters for size, time-series, and bulk samples. Due to the low biomass of the 0.1-µm pore-size samples, multiple 0.1-µm filter samples had to be pooled for proteomics measurements (*i.e.,* three to four 0.1-µm time-points constitute one sample; see Table S3 for the exact list of pooled samples). Additional pooling of the corresponding 0.2-µm filter samples were also combined to serve as a comparison. Furthermore, one pooled sample resulting from a combination of 0.2-µm filters for whom their corresponding 0.1-µm filter samples were insufficient for proteomics, was measured. For metaproteomics of the bulk sample, the leftover 0.1-µm filter from metagenomics was used directly. The time-series measurements were done by taking 2.25 g of the total frozen mass of each 0.2-µm filter pair and pooling the material, resulting in one sample of 0.2-µm filters per time point, each with an approximate weight of 4.5 g. In a similar manner, 4.5 g of the respective 0.1-µm filters were taken, with each generated 0.1-µm sample corresponding to a time-point.

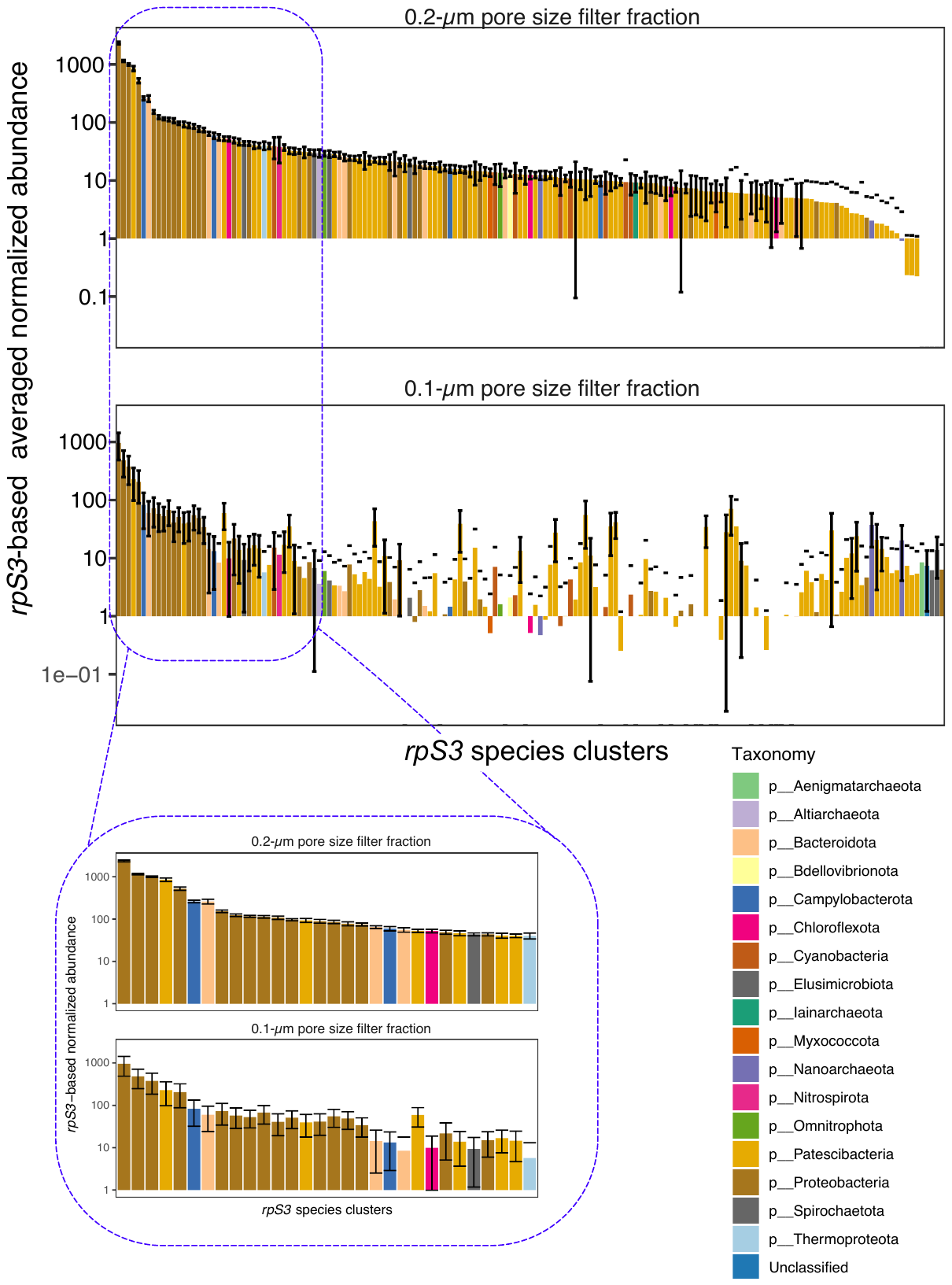

**Supplementary Figure S3. Rank abundances of the WB microbial community.** Species-level normalised rank abundances for the 0.2-µm and 0.1-µm filters. Species are ordered by rank on the 0.2-µm filter, with error bars representing the standard deviation. Abundances are shown in log10-scale and are based on averaged *rpS3* marker gene. The blue square denotes the topmost abundant species in the geyser’s community. The shown color-coded taxonomy classification is based on phylum-level for better visualization.

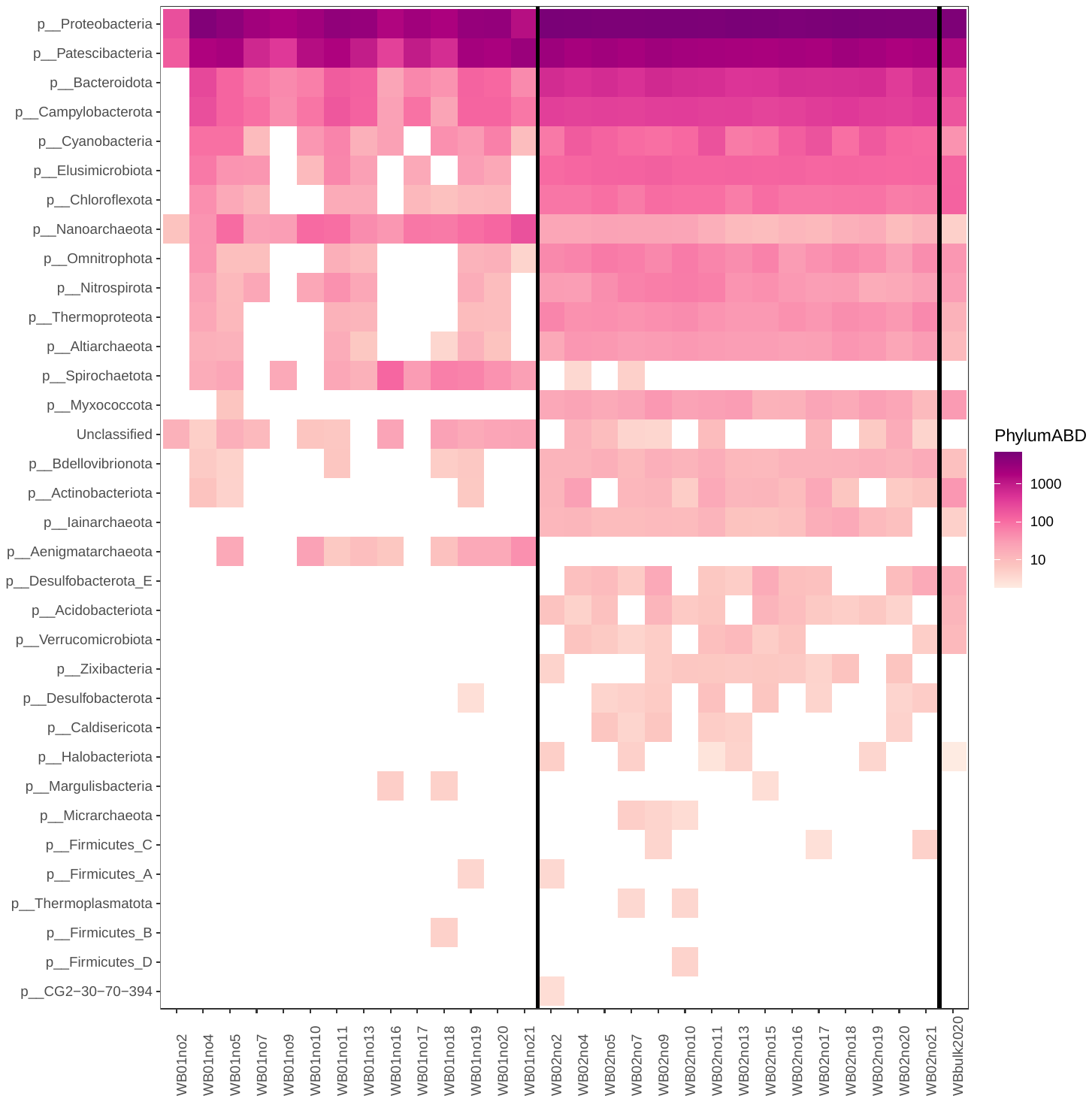

**Supplementary Figure S4. Abundances across time-series of WB microbial community.** Phylum-level normalised abundances of the microbial community of the geyser Wallender Born across all metagenomics time-series samples: 0.2-µm filter (WB02noXX), 0.1-µm filter (WB01noXX), and the 0.1-µm bulk filter sample (WBbulk). All abundances are based on normalized *rpS3* marker gene and shown in log10-scale (Supplementary Table S6).

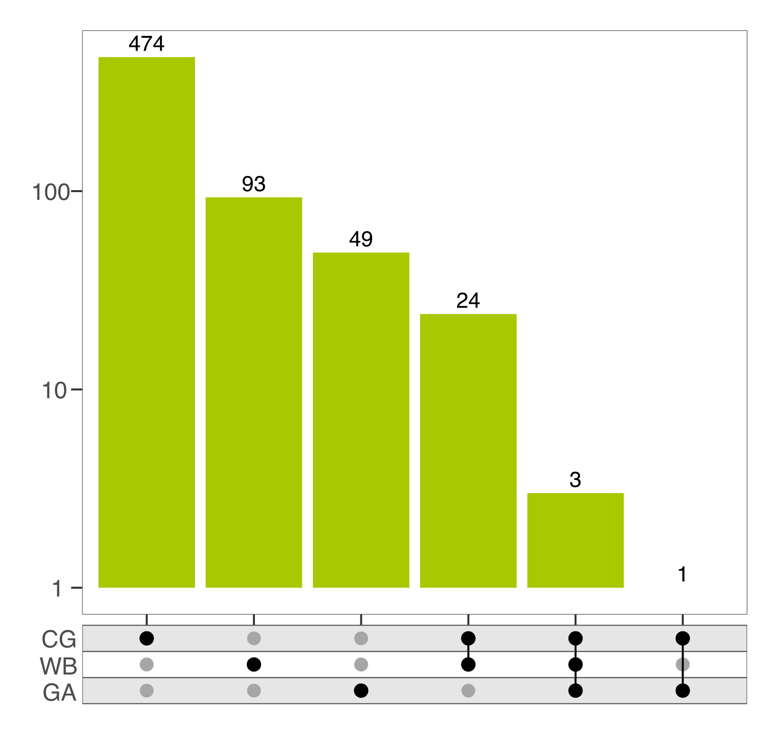

**Supplementary Figure S5. Comparison of the microbial community from cold-water geysers.** Crystal Geyser (CG; Utah, USA), Geyser Andernach (GA; Andernach, Germany) and Geyser Wallender Born (WB; Wallenborn, Germany) are shown in the Y-axis, with those having similar microbial populations connected. X-axis denotes the number of genomes analyzed (FastANI >75% and coverage >50%), as well as the number of shared genomes between sites.

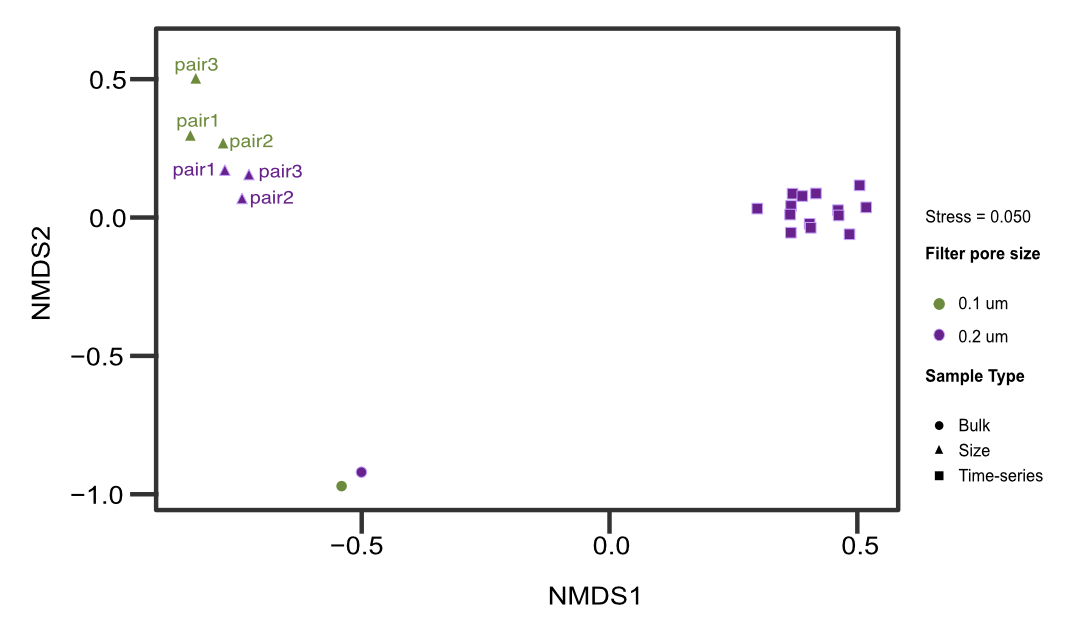

**Supplementary Figure S6. NMDS of metaproteomics samples.** All three sample types analysed in this study are shown to cluster together irrespective of the type of sample in question (time-series, size comparison, and bulk). For the size comparison sample, each corresponding 0.1-/0.2-µm filter pairs are denoted by paired numbering 1, 2, or 3, with green denoting 0.1-µm samples and purple showing the 0.2-µm samples.

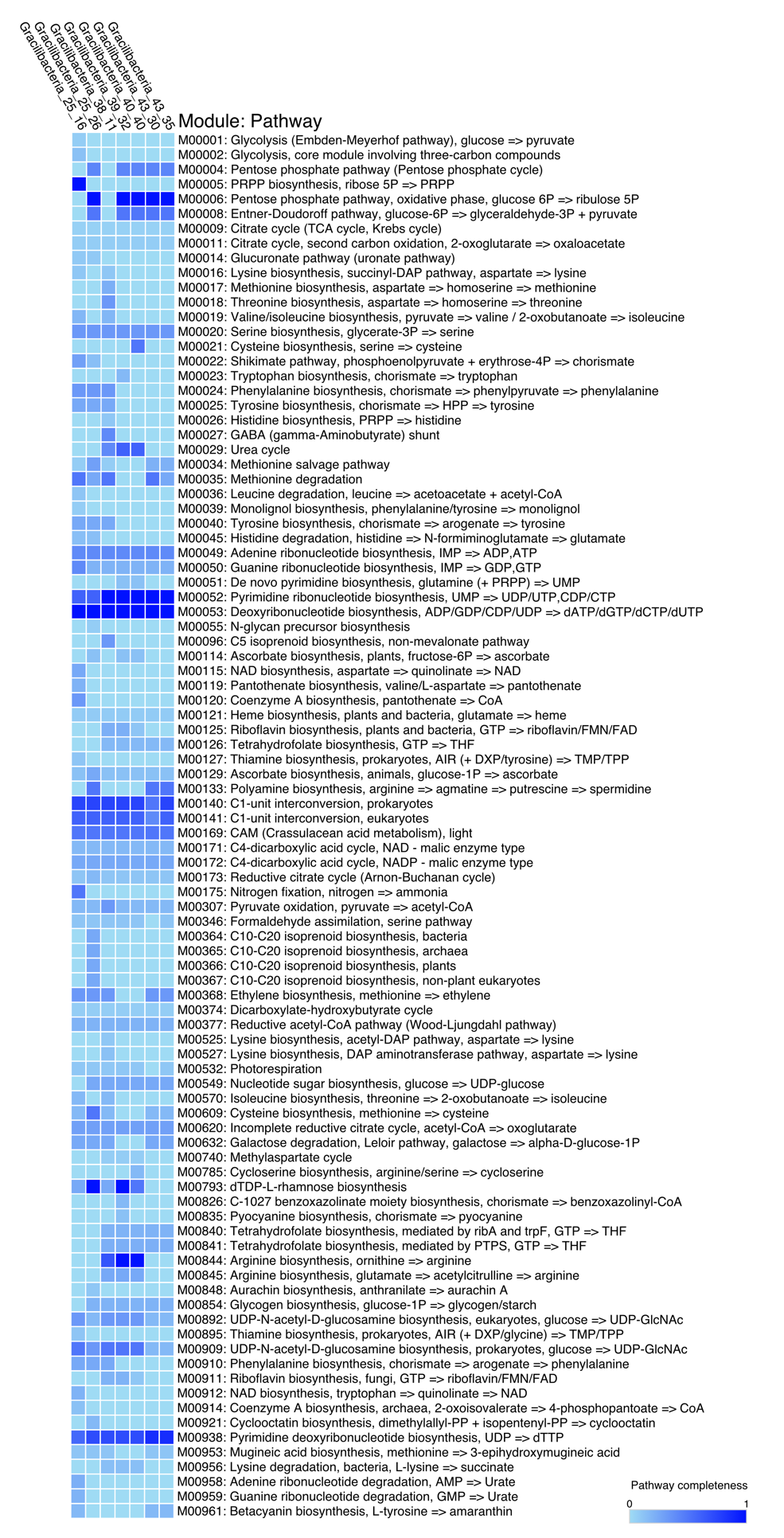

**Supplementary Figure S7. Complete automatic annotation of the Gracilibacteria genomes.** This annotation was performed using the KEGG modules and pathways. The values for the completeness of each pathway can be found in Supplementary table 8.

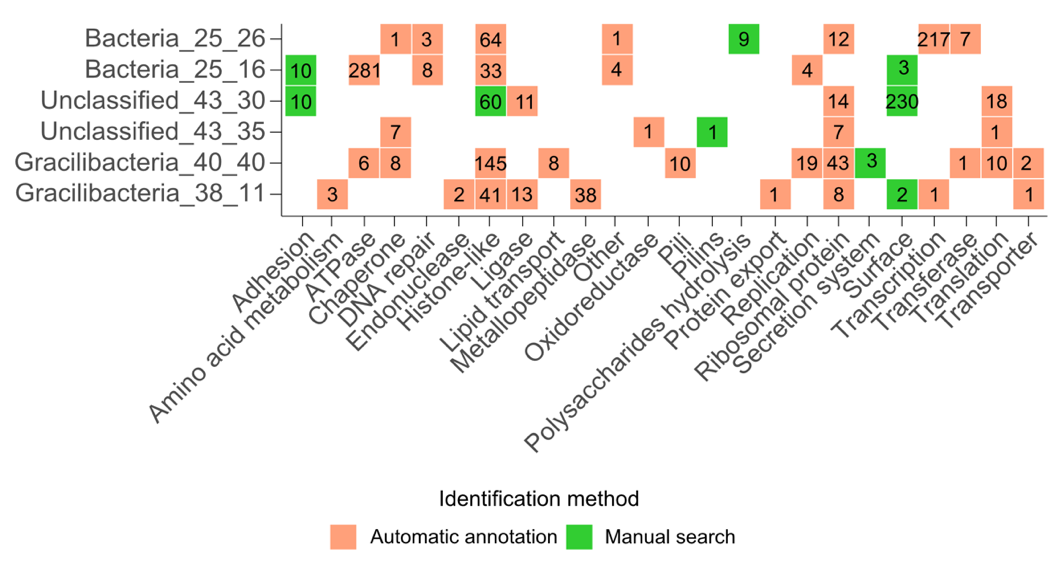

**Supplementary Figure S8. Identified proteins expressed by WB Gracilibacteria.** Proteomic counts of each identified protein, from all samples – time-series, size, and bulk-, were summed up per Gracilibacteria genome. Salmon color denotes proteins that were identified via automatic annotation, while green denotes the protein groups that were identified by manual search of protein domains in public databases.

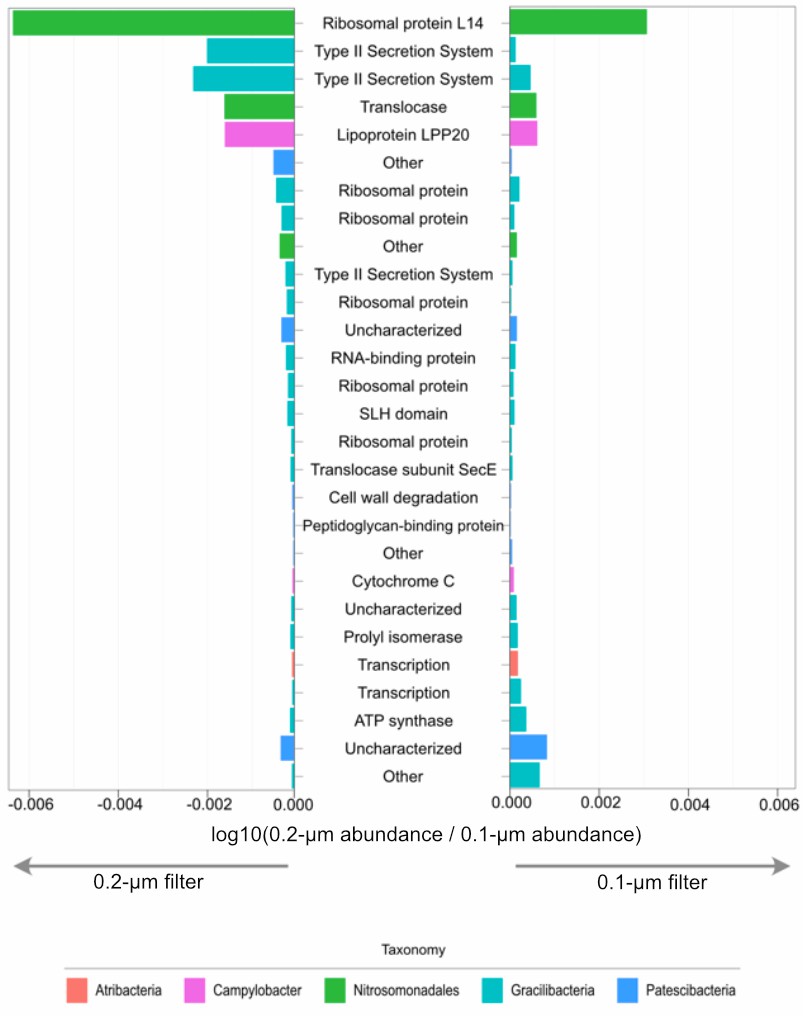

**Supplementary Figure S9. Differential expression of the most abundant bacterial proteins between the 0.2- and 0.1-µm filtered fractions.** Colored bar charts denote the corresponding taxonomy, panel with negative values on the left panel showing the proteins’ abundance on the 0.2-µm filter, while the positive values on the right-side panel denotes the protein expression on the 0.1-µm filter.

### Supplementary Tables

All supplementary tables are provided in the Supplementary_tables.xlsx Excel file, with individual work sheets containing the following tables:

**Table S1.** Filtered volume and cell counts for the WB time-series

**Table S2.** Geochemical, pH, and temperature data obtained for each time point.

**Table S3.** Sample distribution and pooling of metaproteomics samples.

**Table S4.** Coverage and diversity data of raw reads and assembly statistics.

**Table S5.** Normalized rpS3-based abundances across time-series

**Table S6.** Genome information, statistics, and abundances from both the metagenomics and metaproteomics

**Table S7.** Automatic annotation based on KEGG modules for the Gracilibacteria genomes.

**Table S8**. Annotation based on manual search of protein domains.

### Supplementary Files

**File S1**. Metaproteomics database in fasta format. The database is available for the reviewers via the link

<https://nxcl.biologie.uni-due.de/s/wagc5WWP2xAjgzH> and password dEdmMgzs57 and will be get a permanent link on Figshare.

File_S1_Proteomics_database.fasta

**File S2** Metadata of the metaproteomics database (headers). The database is available for the reviewers via the link

<https://nxcl.biologie.uni-due.de/s/wagc5WWP2xAjgzH> and password dEdmMgzs57 dEdmMgzs57 and will be get a permanent link on Figshare.

File_S2_Proteomics_db_metadata.tsv

**File S3.** Phylogenetic placement of Bacteria. Bacteria were taxonomically classified using GTDB-tk, release r207.

File_S3_GTDBtk_Bacteria.tree

**File S4**. Phylogenetic placement of Archaea. Archaea were taxonomically classified using GTDB-tk, release r207.

File_S4_GTDBtk_Archaea.tree
